# Milking it: repeated post-weaning suckling events in Galápagos Sea Lions *Zalophus Wollebaeki*

**DOI:** 10.1101/2025.03.26.645415

**Authors:** Alexandra Childs, Carlina Feldmann, Svenja Stoehr, Rémi Demarthon, Paolo Piedrahita, Sean D. Twiss, Oliver Krüger

**Author notes:** **Corresponding Author:** Alexandra Childs. Department of Animal Behaviour, Faculty of Biology, Bielefeld University, Konsequenz 45, 33501 Bielefeld, Germany (Alexandra Childs).

## Abstract

Milk is not cheap. Mammalian offspring are expected to wean during the early juvenile stages, whether through their own mechanisms or their mothers’ instigation. To not do so would result in a reproductive trade-off for the mother: continued investment in the health and growth of her current offspring, over a shorter birth interval and the possibility of begetting a fitter pup. Here we show for the first time, using 20 years of data, repeated suckling events between female Galápagos Sea Lions (GSL) *Zalophus wollebaeki* and their fully adult biological offspring well beyond the expected age of independence and when the offspring are themselves already reproductively active. This behavior, ‘supersuckling’, suggests that GSL mother-offspring relationships are more complex and longer lasting than previously thought. To our knowledge, this behavior has not been previously documented in any marine mammal species.

## Introduction

At a given point in any mammal’s life, offspring are expected to wean, they cease suckling at their mothers’ teat and achieve nutritional independence by becoming self-sustaining foragers. When this occurs varies greatly both interspecifically (Lee et al. 1991; Derrickson 1992; Pomeroy 2011) and intraspecifically (Trillmich et al. 2014; Eckardt et al. 2016; Matsumoto 2017), depending on their life history, habitat, and food supply. Intraspecific variation has been shown to be influenced by environmental conditions (Georges & Guinet 2000; Trillmich 1990), proximity to resources (Burkanov et al. 2011; Hastings et al. 2021), offspring sex (Lee & Moss 1986), maternal experience and an individual’s position in a social hierarchy (Eckardt et al 2016; Carboni et al. 2022). Amongst marine mammals, pinnipeds employ an extraordinary range of lactation strategies due to their terrestrial or floe ice breeding and aquatic foraging (Trillmich 1996; Pomeroy 2011). Their milk is extremely rich in fat and therefore a costly resource for mothers (Trillmich & Lechner 1986). Eared seals (*Otariidae*) are income breeders employing a prolonged lactation strategy (4 - 36 months, Sepúlveda & Harcourt 2021) where mothers regularly leave in order to forage throughout their offspring’s dependency. Weaning is typically gradual and the literature that addresses this period in *otariids* offer age windows during which weaning should occur (Costa 2001; Sepúlveda & Harcourt 2021).

In *otariids* weaning age is correlated with latitude, with tropical and temperate species attaining independence latest (Trillmich 1996). Nursing beyond what is considered typical weaning age has been documented in several species (e.g. Trites et al. 2006; Lowther & Goldsworthy 2016; Osiecka et al. 2020), highlighting the sometimes extreme variation between individuals in the duration of maternal care. In these previous reports, although duration of observed maternal care was considered unusual relative to species norms, in all cases the durations were considered to be within the limits of the expected transition to independence. However, these studies were also unable to continue observing offspring beyond their second or third year when independence typically occurs, and so could not confirm if these observations were indeed their last suckling bout. Once weaned, juveniles leave their birth site either permanently or until they become reproductively active, and mothers will give birth to their next offspring if they have not already done so (Trillmich & Wolf 2008; Trillmich et al. 2014).

Galápagos Sea Lions (GSL) *Zalophus wollebaeki* are an endangered species endemic to the Galápagos (IUCN: Trillmich 2015), and have suffered a 50% population decline over the last 40 years (Riofrío-Lazo & Páez-Rosas 2023). Females give birth to a single pup on average once every 2 years (Kalberer et al. 2019) and exhibit strong sexual dimorphism from birth (Kraus et al. 2013). As an equatorial species, early life is characterized by one of the longest dependency periods of any *otariid*, and a highly variable weaning age with reports ranging from 1 to as late as 5 years of age (Trillmich & Lechner 1986; Costa 1991; Sepúlveda & Harcourt 2021).

We used 20 years of suckling observations from a colony of GSL on Isla Caamaño and propose an age window during which juvenile GSL are expected to become nutritionally independent. We also describe a previously unknown behavior, where offspring continue to suckle their mothers beyond this window, at regular intervals, throughout their juvenile and adult lives. We investigate the distribution of this behavior within the colony and its relationship with sex and El Niño Oscillations.

## Methods

Data were collected between September 2003 and April 2023 on Isla Caamaño (0°45’S, 90°16’W) during 2 annual field seasons from October-December and February-April that cover the main reproductive period (Pörschmann et al., 2010). The islet is located off the southern coast of Santa Cruz in the centre of the Galápagos archipelago and is home to a breeding colony of GSL that has been intensively studied since 2003 (Wolf & Trillmich, 2007). All individuals that are born on, or regularly use the islet, are marked and assigned a unique identification resulting in a database of extensive individual life history (see Wolf et al., 2005; and Mueller et al., 2011). Pups undergo their first molt at 5-6 months after which they are referred to as ‘juveniles’. Identification rounds were conducted at least twice daily. During each round the ID, location, and behavior (including suckling events) of every individual present on the islet is recorded (Trillmich et al., 2016). Suckling is only recorded when the individual at the teat is clearly suckling, i.e. the head is gently moving back and forth and there is an indentation in the female’s belly caused by the pressure of the suckling individual’s snout, suckling is also frequently accompanied by a distinct slurping noise.

To assess with certainty whether an individual continues to suckle beyond weaning, only offspring with known birth dates who could be assigned to their respective cohort were included in this study. Due to the extended duration of the reproductive period (September - January, Pörschmann et al., 2010), an offspring’s age was assigned according to their reproductive year of birth (season age) rather than the exact day or month. We defined weaning age based on previous juvenile foraging studies (Jeglinski et al. 2012; Piedrahita et al. 2014) and visual inspection of suckling events over time. Offspring who continued to suckle beyond weaning were classed as supersucklers. To determine whether supersuckling is related to alloparenting (Stirling 1975) and milk-stealing (Maniscalco et al. 2007; Reiter et al. 1978), we investigated the occurrence of allonursing within this colony. We inspected the distribution of these supersucklers by their sex, season age and season of birth, using Chi-Squared and Fisher’s Exact tests to assess their relationship with the behavior. We used a Chi-Squared test to examine the influence of El Niño oscillations on the occurrence of supersucklers. Temperature anomaly data was obtained from US National Oceanic and Atmospheric Administration (NOAA 2023) and categorized by El Niño, La Niña and Intermediate conditions according to the Niño Oceanic Index (ONI).

## Results

We found that by 3 seasons of age 90% of offspring had ceased to be observed suckling and the number of last suckling bouts steeply declined between offsprings’ 2^nd^ and 3^rd^ season (Figure 1), which we have interpreted to represent weaning age. We define weaning as an expected window of independence that may occur between 2.5 and 3 seasons of age. Offspring who are observed suckling beyond their 3rd seasons are classed as ‘supersucklers’ and all individuals observed on the islet beyond this window of independence but not seen suckling as ‘non-supersucklers’.

**Figure 1.**
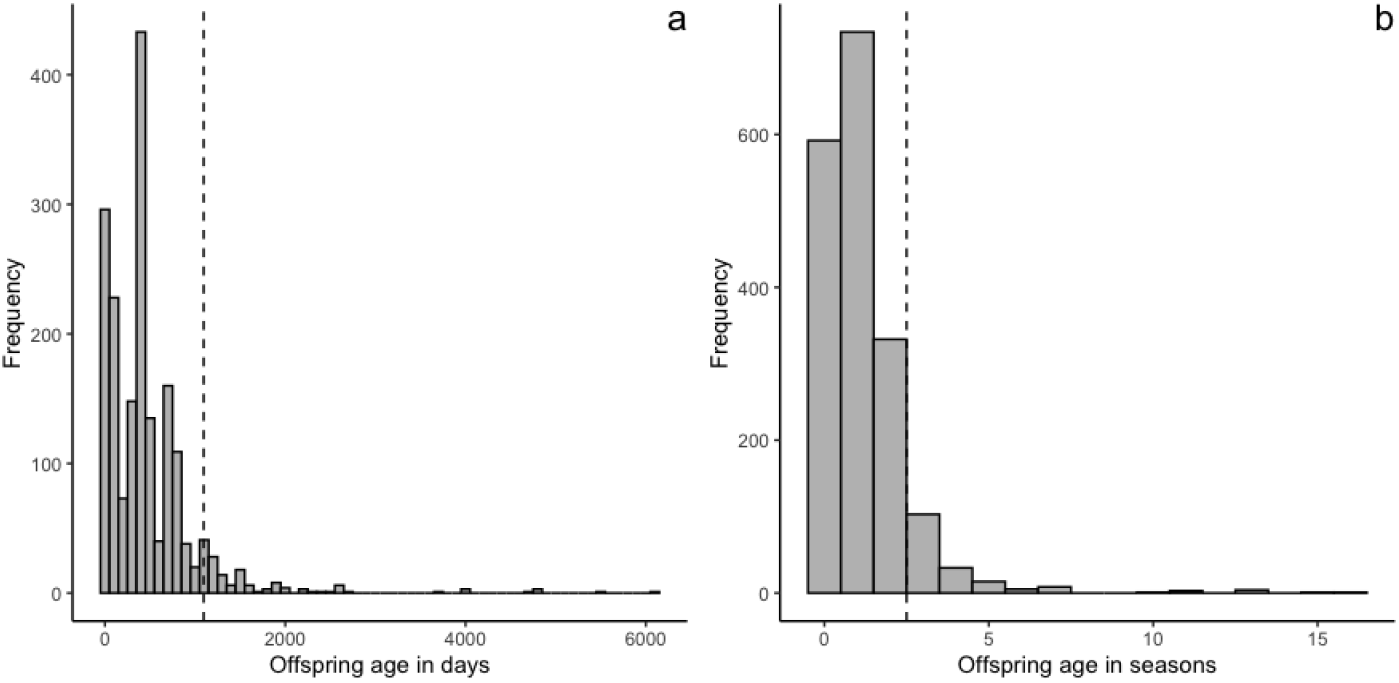
Age of all offspring in **a**) days and **b**) seasons at their last recorded suckling event. Only individuals with a known date of birth are included, *n* = 1832. The dashed line in both **a & b** represents weaning age: **a**) 1095 days, and **b**) 3 seasons where 0 = offspring’s season of birth and weaning is a window between 2.5 and 3 years of age.

**Figure 3.**
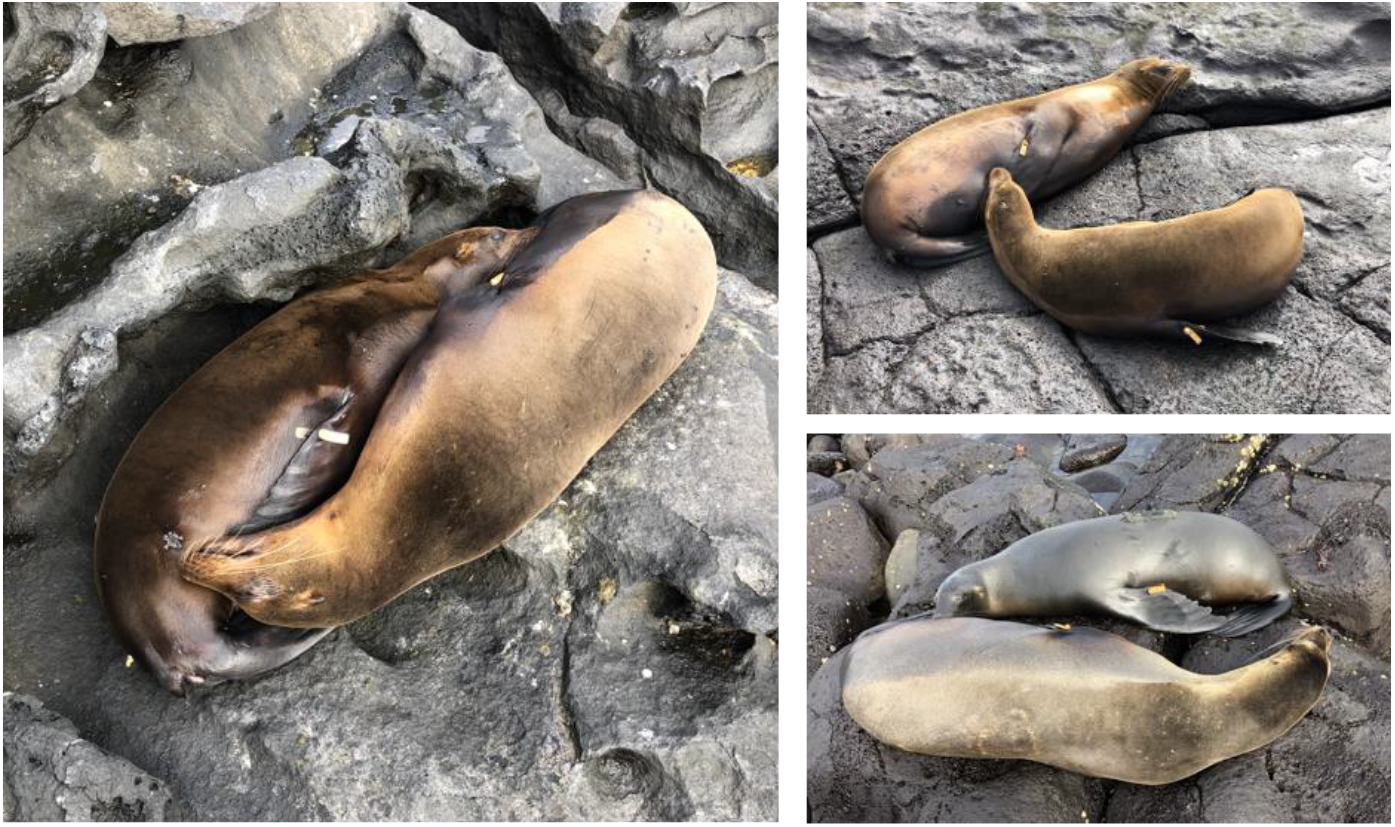
Supersuckling events seen during the reproductive season of 2022, all ages given are those at the time of recording: a) Adult female 444€ on the right, 16 seasons old, suckling her male offspring 414€, 4 seasons old. 414€’s testicles are clearly visible between his hind flippers indicating that he is of, or fast approaching, an age where he could be reproductively active. b) A second supersuckling event with adult female 444€ and her offspring 414€, here it can be seen that 414€ is of a size where we would expect him to be capable of foraging independently. c) Adult female 384€, 17 seasons old, nursing her 3 season old male offspring 502€. Both individuals have be part of an ongoing foraging study using TDRs secured by a neoprene and mesh patch, 384€ has recently lost her patch and 502€ lost his a few weeks after this photograph was taken.

Of the 54,251 suckling events observed on Caamaño islet between 2003-2023, 2145 (4%) were supersuckling. We found that supersuckling was a regular occurrence, observed repeatedly every field season from 2006 onwards (1-10% of all observations). Between 2007-2013, additional identification rounds, with the express purpose of recording suckling events, were conducted for a project on maternal investment. A moderate positive linear relationship (r = 0.56) was found between the number of rounds and supersuckling observations indicating that effort impacts the frequency of observing the behavior but not the likelihood.

The mean age of supersucklers is 1289 days (± 406) whereas the mean age of non-supersucklers is 278 days (± 276), with the suckling population mean at 346 days (± 321). Supersucklers are unevenly distributed according to age with 95% occurring between 3 and 6 seasons of age (Table 1). GSL are considered adults and therefore sexually mature, once they reach 5 years of age (Trillmich et al. 2014; Krüger et al. 2021). As pups have only been tagged since 2003 the maximum possible age for a supersuckler would be 20. The maximum recorded age of a supersuckler in this study was 16 seasons of age and the presence of multiple mature supersucklers shows that for some individuals this behavior is maintained well into adulthood. As the project continues, the number of mature supersucklers is likely to continue to increase. Of the offspring old enough to be classified (*n* = 1736), 186 are supersucklers (11%), 38 (20%) of which continued to suckle into their adulthood and accounted for 11% of all supersuckling events recorded.

**Table 1.**
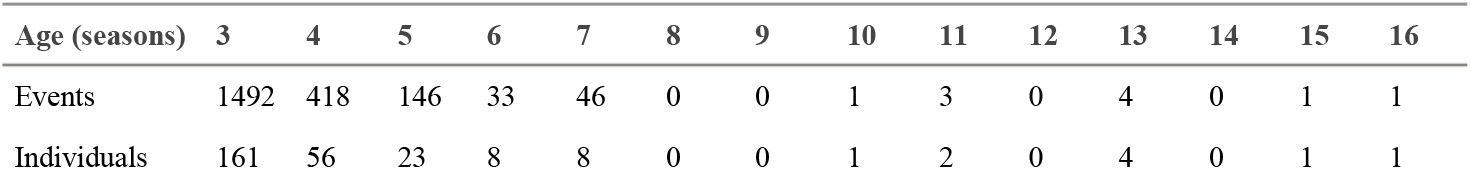
Distribution of observed supersuckling events and the number of unique supersucklers according to age in seasons.

For the purposes of this study, 3 video recordings of supersuckling events were made during the Spring season of 2024 (see supplementary material). Video 1 shows a female nursing her 4.5 year old offspring on two separate occasions, the second of which she is clearly expressing milk from the teat her offspring has just left. Video 2 shows a second mother offspring pair of the same age where the size of the offspring is almost equal to that of its adult mother. Figure 2 further illustrates the similarities in size between mother and offspring and their approaching maturity. All of these observations are of identified individuals with exact dates of birth. Video 3 is a chance observation of what is highly likely to be a supersuckling incident at San Cristobal Malecón. The offspring here is not much smaller than the nursing female and is therefore likely to be at least 3 seasons or older, however we are unable to confirm this.

In a highly unusual occurrence of supersuckling, the authors witnessed a tagged 6-year-old adult female (A) supersuckling another unidentified adult female (B) whilst A was simultaneously being suckled by her first pup (C). Video 4 shows this trio just after they have finished suckling but are still in contact with each other. This suckling ‘train’ occurred on multiple occasions over two separate field seasons during the 2022 reproductive year. Female B was always at the head of this ‘train’ with female A supersuckling in second, and her pup C suckling at the tail. This order never changed when the train was in progress and although female A and B, and A and C were seen suckling in separate pairs, no allonursing was observed between C and B or any other unidentifiable female. We identified all allonursing events within the data set to determine whether supersuckling is an extension of alloparenting and milk-stealing. As we could not confirm kinship of unidentified nursing females, all suckling events without a positive ID for the nursing female or biological mother were eliminated. We define alloparenting as offspring having been observed suckling both biological mother and a non-filial female more than once, and all one-off allonursing events as milk-stealing. We found 123 instances of milk stealing of which 6 (5%) were supersuckling, and 275 allonursing events (2% of all suckling events). Of the 1832 suckling offspring observed, 29 benefited from alloparenting and approximately 1:6 became supersucklers.

The proportion of suckling events is evenly distributed between the sexes until 5 seasons of age when the proportion of males became much higher than the females before reversing at 7 seasons. Due to the small number of adult supersuckling events (*n* = 235), proportions beyond 5 seasons should be treated with caution. Overall, equal proportions of females (51%) and males (49%) have been recorded supersuckling. Female supersuckler mean age (609 days ± 6.34) is slightly higher than males (590 days ± 6.75), however, a chi-squared test found no significant relationship between sex and the act of supersuckling (*p =* 0.174) or being a supersuckler (*p =* 0.090).

Supersucklers have been born every reproductive season from 2003 - 2019, offspring born afterwards were not old enough at the time of this study to be classified. We found a significant relationship (*p <* 0.001) between reproductive season of birth and the probability of offspring becoming supersucklers. To determine if this relationship could be explained by environmental oscillation events, we used monthly temperature anomalies (ONI, NOAA 2023) and a Chi-squared test, but found no relationship (*p =* 0.594). However, there is a significant relationship (*p* < 2.2×10^−16^) between the type of suckling event and the type of oscillation (ONI, NOAA 2023). Further investigation of the deviation from the expected number of observations showed that supersuckling is much more likely to occur under La Niña conditions than any other oscillation type.

## Discussion

Weaning age has been historically challenging to pinpoint in GSL with literature suggesting that independence may occur from as early as 1 to as late as 5 years old (see: Trillmich & Lechner 1986; Trillmich et al. 2014; Krüger et al. 2021). Here we show that juvenile GSL are most likely to wean by their third reproductive season or between 2.5 and 3 years of age (Figure 1). What is highly unusual is the number of offspring who continue to suckle beyond this window as both young and mature adults. Being nutritionally dependent over multiple years is common in other large mammals such as orca, *Orcinus orca* (Foster et al. 2012), orangutans, *Pongo pygmaeus wurmbii* (van Noordvijk et al. 2013), and elephants, *Loxodonta africana* (Lee & Moss 1986), however these are all long lived species (maximum age of 40-120 years). GLS live comparatively short lives reaching a maximum age of 21 years (Krüger et al. 2021), meaning offspring continuing to suckle beyond the weaning window accounts for 1/5 to 1/4 of their lives. In human terms, if the average lifespan is 80 years, this is the equivalent of a 16-20 year old child still breast-feeding.

That so many supersucklers were sexually mature at the time of observation is unprecedented. It is clear from observations such as Video 4 that many, if not all, of these individuals are reproductively active. Supersuckling may increase a mature offspring’s reproductive success, which could be considered as delayed generational success for the mother, offsetting any energetic or reproductive cost in allowing her mature offspring to suckle her. Given their age and size, it is highly improbable that these supersucklers are not already foraging to the extent that they could be considered nutritionally independent (Jeglinski et al. 2012). In the majority of supersuckling events that we have witnessed, offspring appear both large and healthy enough to forage independently (Videos 1-2, Figure 2).

GLS are sexually size dimorphic with adult males weighing up to 158kg and females averaging 60kg (Trillmich et al. 2014). As previous work has indicated that males demand and receive a higher level of investment from their mothers (Kraus et al. 2013; Piedrahita et al. 2014), we were surprised to find that offspring sex does not seem to play a role in this behavior. This is possibly because as male offspring reach sub-adulthood, dominant males within the colony perceive them as a threat and chase them off, separating many from their mothers and preventing them from supersuckling. However, this does not explain why males are still almost equally likely to be seen supersuckling as females.

Unlike in some phocid seal species (Cassini 1999), lactation does not delay oestrus in GSL and females become receptive within a few weeks of parturition (Heath 1989). Consequently, females remain in a permanent cycle of pregnancy and lactation throughout their reproductive lives (Trillmich et al. 2014; Krüger et al. 2021). Although we cannot assume that this is the case for all supersuckling events, it seems reasonable to infer that females nursing a supersuckling offspring would be expressing milk due to their cyclical reproductive state after primiparity. Likewise, it is unlikely offspring would repeatedly suckle beyond the weaning window without receiving some form of nutritional compensation, even if the benefits are minimal. As milk is such a costly resource (Trillmich & Lechner 1986), it is important that further research is undertaken to understand who the driver in this mother-offspring relationship is, what the circumstances are that allow this behavior to come about, and the subsequent costs and/or benefits to each.

Female GSL give birth every 2 years (Kalberer et al. 2019), although intraspecific variation is common and many females give birth whilst their previous offspring is still dependent. Females have occasionally been observed nursing two dependent offspring simultaneously but the majority of mothers reject one shortly after birth (Trillmich & Wolf 2008). This would imply that allowing an offspring to become a supersuckler would represent a reproductive trade-off for the mother. As GSL females are highly protective of their offspring and aggressively dissuade any non-filial attempts to suckle (Trillmich & Wolf 2008), it is not surprising that so few allonursing and milk-stealing events were recorded. The high ratio of alloparented offspring becoming supersucklers indicates that there is a relationship between the two but given the total number of alloparented offspring (29) compared to the number of supersucklers (186), the two should be considered distinct from each other. That only 5% of milk stealing events were also supersuckling indicates that this behaviour is different from the post-weaning milk-stealing that has been observed in male northern elephant seals *Mirounga angustirostris* (Reiter et al. 1978). The extremely rare occurrence of allonursing and milk-stealing in this colony also reinforces the widely accepted assumption that milk is costly to the mother and therefore a precious resource to be invested wisely (Trivers 1974; Boness & Bowen 1996). Observations like Video 4 indicate that this investment ends with a female’s own offspring regardless of the proximity of their kinship. It does not, however, explain why mothers continue to invest milk in offspring who are independent foragers and reproductively active. Lifetime mother-pup bonding without suckling has previously been observed in captive California sea lions *Zalophus californianus* (Hanggi & Schusterman 1990) which could suggest a social basis for these post-weaning interactions as has been found in orca (Weiss et al. 2023) and chimpanzees, *Pan troglodytes verus* (Samuni et al. 2020).

As in other *otariids*, individual variation in weaning age may occur due to resource proximity, abundance, or the presence of a rival sibling (Trillmich & Wolf 2008; Burkanov et al. 2011). The females of the Caamaño islet population have diverse, but stable foraging strategies that target different prey species in different locations, to which they maintain a high level of fidelity (Schwarz et al. 2022; Stoehr et al. *unpublished*). We would therefore not expect proximity to foraging grounds to impact weaning age as has been shown in Steller sea lions, *Eumetopias jubatus* (Hastings et al 2021), but rather food abundance. GSL are exposed to extreme changes in environmental conditions caused by the El Niño oscillations (Trillmich et al. 2014). Although mothers have been shown to buffer against this during an offspring’s dependency (Kraus et al. 2013), we anticipated that during El Niño conditions, supersuckling would be more common as mother-offspring pairs delayed weaning until more favorable conditions return. Contrary to expectations, we found that supersuckling was more prevalent during La Niña conditions.

That supersuckling has been observed every field season since 2006 indicates that it is neither rare nor unusual, and the opportunistic mother-offspring observation on San Cristobal (Video 3) suggests that the behavior is likely present in other colonies. It is highly unusual for such a short-lived species to continue to suckle to such an advanced age and as far as the authors are aware, this is the first detailed account of supersuckling in GSL. It is also the first description of repeat suckling events by sexually mature offspring in any mammalian species. The overall population of GSL in the archipelago has declined by more than 50% in the last 40 years (Riofrío-Lazo & Páez-Rosas 2023) and the Caamaño colony has been in a steep decline since 2014 (*unpublished data*). Supersuckling is an example of how much is still left to be understood about GSL and highlights the importance of long-term studies of individuals within a population.

## Supporting information

html code output

## Acknowledgements

We are grateful to all past PI’s, PhDs and field assistants whose tireless work in the field has made this study possible. This publication is contribution number 2659 of the Charles Darwin Foundation for the Galápagos Islands. Research on live animals followed guidelines of the American Society of Mammalogists (Sikes & Gannon 2011). All animal handling and experimentation were performed under Galápagos National Park scientific permits PC-83-21, PC-83-22, and PC-83-23 and made possible by the logistical support of the Charles Darwin Foundation. This work was supported by the German Research Foundation (DFG) via the collaborative research centre CRC TRR 212, project number 316099922, project B07, and Bielefeld University.

## Declaration of interests

The authors declare no conflict of interest.

## Notes

### Competing Interest Statement

The authors have declared no competing interest.

