## Supplementary material for "Milking it: repeated post-weaning suckling events in Galápagos Sea Lions *Zalophus Wollebaeki*": html code output: Code_Supersuckling_BehaviouralNote_Submitted.html

Super Suckler Behavioural Note


### Super Suckler Behavioural Note

#### 2024-05-24

#### Supersuckling definition

… loading in the data set with one row per observed suckling

… loading in growth data set which has information on age when first
observed

… loading in data set that has all observations to identify
individuals that were observed after 3 seasons old

… doing some data cleaning (remove implasuible values, assume birth
dates of all animals born before 06/2003 to be unknown)

… creating new category in `offspring_exact_birth`: not
only exactly(1) and weakly(2) known (+-2 weeks) dates of birth are of
interest, but also roughly(3) known dates if the pup was estimated to be
less than 6 months old when first found.

```
## 
##    0    1    2    3 
##  807 1506  440  243
```

… creating new variables:

- `season` is the season in which the observation was
  made: a count that starts with season 1 (8/2000-7/2001), up to season 24
  (8/2023-7/2024)
- `offspring_sob` is the season in which the observed
  offspring was born, `NA` if date of birth not known
- `offspring_age_seasons` which gives the age of the
  offspring measured in seasons
- `supersuckling` is `TRUE` if the offspring
  was at least 3 seasons old when observed suckling

Defining **what is considered supersuckling** (observed
suckling after the pup is expected to be weaned off) is of course a
researcher’s choice. Using the age in seasons instead of years seems a
bit less arbitrary:

This plot shows all observed suckling events. The events defined as
supersuckling (**suckling when at least 3 seasons old**)
are colored blue. The dashed horizontal line indicates an alternative
criterion for defining supersuckling: when the offspring is at least
three years old.

… defining age of independence which will be the cut off for whether
an offspring is a supersuckler or not

##### Histogram with all sightings

one plot to show the age in days (50 day bins) and one plot to show
the age in seasons (1 bin = 1 reproductive year). This makes it easier
to see the hypersucklers (i.e. mature adults) and also sea lions don’t
celebrate their birthday so seasons is more appropriate for observing
when dependents stop suckling

```
## # A tibble: 1 × 1
##   distinct_offspring_IDs
##                    <int>
## 1                   1832
```

Also the same hisograms but only with the age when they were last
seen suckling:

###### Descriptive numbers for the text

… counting the number of super suckling events (n = 2145) number of
super suckling individuals = 186 (see 4 blocks down)

```
## 
## FALSE  TRUE  <NA> 
## 46174  2145  5932
```

… counting the events for each reproductive year

```
## # A tibble: 20 × 6
##    season supersucklingevents nonsusuevents   NAs  from    to
##     <dbl>               <int>         <int> <int> <dbl> <dbl>
##  1      4                   0           986   753  2003  2004
##  2      5                   0          1777   345  2004  2005
##  3      6                   0          1304   191  2005  2006
##  4      7                  39          2502  1025  2006  2007
##  5      8                 259          4553   758  2007  2008
##  6      9                  82          3926   281  2008  2009
##  7     10                 137          2792   333  2009  2010
##  8     11                 190          3343   169  2010  2011
##  9     12                 556          7080   468  2011  2012
## 10     13                 255          5512   418  2012  2013
## 11     14                 241          3590   218  2013  2014
## 12     15                 122          1823   184  2014  2015
## 13     16                  63          1568   194  2015  2016
## 14     17                   6           592    77  2016  2017
## 15     18                  33          1428    54  2017  2018
## 16     19                  31          1167   101  2018  2019
## 17     20                  11           937    63  2019  2020
## 18     21                   6            30    22  2021  2022
## 19     22                  47           674   137  2021  2022
## 20     23                  67           590   141  2022  2023
```

… counting the number of mega sucklers = 4 seasons or older …
counting the number of sexually mature/hyper sucklers = 5 seasons or
older

```
## [1] "number of sightings of individuals who are 4 seasons or older"
```

```
## [1] 653
```

```
## [1] "number of unique IDs seen at 4 seasons or more"
```

```
## [1] 74
```

```
## [1] "number of sightings of individuals who are 5 seasons or older"
```

```
## [1] 235
```

```
## [1] "number of unique IDs seen at 5 seasons or more"
```

```
## [1] 38
```

###### Does the number of rounds impact likelihood of observing supersuckling?

###### correlation test

```
## 
## Call:
## lm(formula = supersuckling_count ~ rounds_suckling, data = sucklingrounds)
## 
## Residuals:
##     Min      1Q  Median      3Q     Max 
## -4.3260 -1.3642 -0.3642  0.6358 10.6867 
## 
## Coefficients:
##                 Estimate Std. Error t value Pr(>|t|)    
## (Intercept)     -1.59764    0.12002  -13.31   <2e-16 ***
## rounds_suckling  0.98728    0.03722   26.53   <2e-16 ***
## ---
## Signif. codes:  0 '***' 0.001 '**' 0.01 '*' 0.05 '.' 0.1 ' ' 1
## 
## Residual standard error: 1.65 on 1545 degrees of freedom
## Multiple R-squared:  0.3129, Adjusted R-squared:  0.3125 
## F-statistic: 703.7 on 1 and 1545 DF,  p-value: < 2.2e-16
```

```
## [1] 716
```

#### Is this a random subset?

Checking distribution by 1. sex 2. birth

… creating a new df with all observations and essential offspring
information … ‘supersuckling\_event’ tells if the associated suckling
event was supersuckling, ‘supersuckler\_ID’ tells if associated ID has
ever been seen super suckling (TRUE)

```
## [1] "Number of unique IDs who do (TRUE) or do not super suckle:"
```

```
## 
## FALSE  TRUE  <NA> 
##  1646   186   290
```

```
## [1] "Population Sex Distribution:"
```

```
## 
##    Female      Male 
## 0.4900971 0.5099029
```

```
## [1] "Super Sucklers Sex Distribution:"
```

```
## 
##    Female      Male 
## 0.5083481 0.4916519
```

```
## [1] "Population Age Summary:"
```

```
##      Min.   1st Qu.    Median      Mean   3rd Qu.      Max.        SD 
##    0.0000   55.0000  337.0000  346.1323  424.0000 6079.0000  321.6823
```

```
## [1] "Supersucklers Age Summary:"
```

```
##      Min.   1st Qu.    Median      Mean   3rd Qu.      Max.        SD 
##  798.0000 1076.0000 1126.0000 1289.1883 1449.0000 6079.0000  405.8987
```

```
## [1] "Non-Supersucklers Age Summary:"
```

```
##     Min.  1st Qu.   Median     Mean  3rd Qu.     Max.       SD 
##   0.0000  44.0000 317.0000 278.3805 389.0000 981.0000 226.0276
```

```
## # A tibble: 2 × 3
##   offspring_sex average_age_at_sighting_days standard_error
##   <chr>                                <dbl>          <dbl>
## 1 Female                                609.           6.34
## 2 Male                                  590.           6.75
```

##### Visualisation and testing

###### Sex

Same plot for distinct individuals instead of total number of
sightings and also including the number of individuals as size of the
points:

… testing if there is a relationship between sex and supersuckling
events

```
## 
##  Chi-squared test for given probabilities
## 
## data:  sex_event_chisquared_table
## X-squared = 1.8503, df = 1, p-value = 0.1737
```

… testing if there is a relationship between sex and being a
supersuckler

```
## 
##  Chi-squared test for given probabilities
## 
## data:  sex_ID_chisquared_table
## X-squared = 2.8383, df = 1, p-value = 0.09204
```

the proportion of males and females seen suckling seems to be evenly
distributed until 4 years of age after which it goes mad but this is
likely due to the decrease in sample size after 5 years of age.

The results of the chi-squared test sow that sex is not significantly
related to either the act of supersuckling (p = 0.1737) or being a
supersuckler (p = 0.09204)

###### Age

```
## # A tibble: 13 × 2
##    offspring_age_seasons true_count
##                    <dbl>      <int>
##  1                     0          0
##  2                     1          0
##  3                     2          0
##  4                     3       1492
##  5                     4        418
##  6                     5        146
##  7                     6         33
##  8                     7         46
##  9                    10          1
## 10                    11          3
## 11                    13          4
## 12                    15          1
## 13                    16          1
```

```
## # A tibble: 10 × 2
##    offspring_age_seasons unique_offspring_count
##                    <dbl>                  <int>
##  1                     3                    161
##  2                     4                     56
##  3                     5                     23
##  4                     6                      8
##  5                     7                      8
##  6                    10                      1
##  7                    11                      3
##  8                    13                      4
##  9                    15                      1
## 10                    16                      1
```

```
## 
## Fisher test for relationship between reproductive year and being a supersuckler:
```

```
## 
##  Fisher's Exact Test for Count Data with simulated p-value (based on
##  2000 replicates)
## 
## data:  as.matrix(ss_age_sex_distribution_group_year)[, 2:3]
## p-value = 0.0004998
## alternative hypothesis: two.sided
```

```
## # A tibble: 2 × 2
##   supersuckler_ID median_reproductive_year
##   <lgl>                              <dbl>
## 1 FALSE                               2010
## 2 TRUE                                2008
```

The plot shows that supersucklers have been born every year since the
start of the project. The last 3 years of data were removed for this and
the following test because individuals born in these reproductive years
are not yet old enough for us to know if they will exhibit this
behaviour. The Fisher test shows that there is a significant
relationship (p = 4.9975012^{-4}) between reproductive year of birth and
being a supersuckler. Perhaps this relationship could be explained by
the El Nino oscillations…

##### El Niño/La Nińa

further testing to see if more supersucklers are born in El Nino
years importing climate data from NOAA (https://origin.cpc.ncep.noaa.gov/products/analysis\_monitoring/ensostuff/detrend.nino34.ascii.txt)
adding it to ss\_age\_sex\_distribution and ss\_age\_sex\_distribution\_group
so that we can see if there is a relationship with oscillation class
visualise data - cannot visualise it very well, so will rely on test
results we will first test if there is a relationship between being a
supersuckler and the oscillation type during the month of their birth
then we will test if there is a relationship between supersuckling
events and the oscillation type

```
## 
##  Chi-squared test for relationship between being a supersuckler and oscillation type during their birth month:
```

```
## 
##  Pearson's Chi-squared test
## 
## data:  table(ss_age_sex_distribution_group$supersuckler_ID, ss_age_sex_distribution_group$category)
## X-squared = 1.3505, df = 2, p-value = 0.509
```

```
## 
##  Chi-squared test for relationship between supersuckling being observed and oscillation type during observation:
```

```
## 
##  Pearson's Chi-squared test
## 
## data:  table(ss_age_sex_distribution$supersuckling_event, ss_age_sex_distribution$category)
## X-squared = 94.838, df = 2, p-value < 2.2e-16
```

```
## Deviation from expected counts for supersuckling events and oscillation type:
```

```
##        
##            El Niño Intermediate    La Niña
##   FALSE  144.24736     47.43902 -191.68638
##   TRUE  -144.24736    -47.43902  191.68638
```

Plots are very unclear. The results of the chi-squared test indicate
that there is no relationship (p = 0.509022) between an individual
becoming a supersuckler based on the oscillation status of its month of
birth. However there is a significant relationship (p<
`r`ssEvent\_chisq$p.value`) between the kind of suckling event
and the type of oscillation. Investigating the deviation from the
expected number of observations we can see that super suckling is more
likely to occur during La Nina.

###### Same time series plots as for birth month also for event month:

#### Alloparenting and milk stealing

checking these behaviours relationship with each other

including only individuals with a known dob as we cannot know if they
are supersucklers without this information

The following tables allow extracting any information relevant to
describe the prevalence of supersuckling and superweaning
(i.e. allosuckling and/or milk stealing). All events and all individuals
are classified as either supersuckling or not and either superweaning or
not. If the offspring’s birth date isn’t known or after 2019,
supersuckling is NA. If the offspring’s birth mother ID or the ID of the
suckled female isn’t known, superweaning is NA.

Furthermore, superweaning can be either milk stealing (if offspring
was seen suckling from an identified female other than its birth mother
only once) or alloparenting (if offspring was seen suckling from another
identified female multiple times).

```
## 
##  Numbers of suckling events who were supersuckling or allonursing (milk stealing or alloparenting)
```

```
##              superweaning
## supersuckling FALSE  TRUE  <NA>   Sum
##         FALSE 15924   359 28778 45061
##         TRUE    794     6  1345  2145
##         <NA>   2103    33  4909  7045
##         Sum   18821   398 35032 54251
```

```
##              alloparenting
## supersuckling FALSE  TRUE  <NA>   Sum
##         FALSE 16024   259 28778 45061
##         TRUE    800     0  1345  2145
##         <NA>   2120    16  4909  7045
##         Sum   18944   275 35032 54251
```

```
##              milk_stealing
## supersuckling FALSE  TRUE  <NA>   Sum
##         FALSE 16183   100 28778 45061
##         TRUE    794     6  1345  2145
##         <NA>   2119    17  4909  7045
##         Sum   19096   123 35032 54251
```

```
## 
##  Numbers of pups who were super sucklers or super weaners (milk stealers or alloparented)
```

```
##             superweaner
## supersuckler FALSE TRUE <NA>  Sum
##        FALSE   523   89  938 1550
##        TRUE     54   22  110  186
##        <NA>    129   19  238  386
##        Sum     706  130 1286 2122
```

```
##             alloparented
## supersuckler FALSE TRUE <NA>  Sum
##        FALSE   590   22  938 1550
##        TRUE     71    5  110  186
##        <NA>    146    2  238  386
##        Sum     807   29 1286 2122
```

```
##             milk_stealer
## supersuckler FALSE TRUE <NA>  Sum
##        FALSE   540   72  938 1550
##        TRUE     58   18  110  186
##        <NA>    131   17  238  386
##        Sum     729  107 1286 2122
```

..extracting some relevant numbers from the tables

```
## [1] "Instances of milk stealing:  123"
```

```
## [1] "Instances of alloparenting:  275"
```

```
## [1] "Instances of suckling with identified suckled female and birth mother:  19219"
```

```
## [1] "Proportion of allonursing of all suckling events"
```

```
## [1] "with identified suckled female and birth mother:  0.0207086737083095"
```

```
## [1] "Supersuckling events out of all allonursing events:  0"
```

```
## [1] "Supersuckling events out of all milk stealing events:  6"
```

```
## [1] "Number of suckling offspring observed:  2122"
```

```
## [1] "Number of suckling offspring observed with known birth mother:  836"
```

```
## [1] "Number of offspring who were alloparented:  29"
```

```
## [1] "Number of offspring who were alloparented and became supersucklers:  5"
```

```
## [1] "Number of offspring who were milk stealers:  107"
```

```
## [1] "Number of offspring who were milk stealers or alloparented:  130"
```

```
## [1] "Number of suckling offspring observed with known age and born before 2020 "
```

```
## [1] "(potential identifiable super sucklers):  1736"
```

```
## [1] "Number of offspring who were super sucklers:  186"
```

```
## [1] "Proportion of offspring who became super sucklers:  0.107142857142857"
```
